# MitoPlex: A Targeted Multiple Reaction Monitoring Assay for Quantification of a Curated Set of Mitochondrial Proteins

**DOI:** 10.1101/820167

**Authors:** Aleksandr B. Stotland, Weston Spivia, Amanda Orosco, Allen M. Andres, Roberta A. Gottlieb, Jennifer E. Van Eyk, Sarah J. Parker

**Author notes:** These authors contributed equally: Jennifer Van Eyk, Ph.D. and Roberta A. Gottlieb M.D. **Corresponding author:** Sarah J Parker, Ph.D. Lead contact: Aleksandr B. Stotland, Ph.D.

## Abstract

Mitochondria are the major source of cellular energy (ATP), as well as critical mediators of widespread functions such as cellular redox balance, apoptosis, and metabolic flux. Methods to quantify mitochondrial content are limited to low throughput immunoassays, measurement of mitochondrial DNA, or relative quantification by untargeted mass spectrometry. Here, we present a high throughput, reproducible and quantitative mass spectrometry multiple reaction monitoring based assay of 37 proteins critical to central carbon chain metabolism and overall mitochondrial function termed ‘MitoPlex’. We coupled this protein multiplex with a parallel analysis of the central carbon chain metabolites (218 metabolite assay) extracted in tandem from the same sample, be it cells or tissue. In tests of its biological applicability in cells and tissues, ‘MitoPlex plus metabolites’ indicated profound effects of HMG-CoA Reductase inhibition (e.g., statin treatment) on mitochondria of i) differentiating C2C12 skeletal myoblasts, as well as a clear opposite trend of statins to promote mitochondrial protein expression and metabolism in heart and liver, while suppressing mitochondrial protein and ii) aspects of metabolism in the skeletal muscle obtained from C57Bl6 mice. Our results not only reveal new insights into the metabolic effect of statins in skeletal muscle, but present a new high throughput, reliable MS-based tool to study mitochondrial dynamics in both cell culture and in vivo models.

## Introduction

Central carbon metabolism is a highly conserved enzyme-mediated network that converts carbon into energy, comprised of sub-pathways such as the oxidative tri-carboxylic acid (TCA cycle), glycolysis, gluconeogenesis and pentose phosphate pathway (Spinelli and Haigis, 2018). Mitochondria are the primary metabolic organelles in the cell where these processes take place (Pagliarini and Rutter, 2013), and therefore play a pivotal role in mediating cellular homeostasis by not only maintaining the oxidative TCA cycle (Martinez-Reyes et al., 2016) and fuel utilization (glycolysis and fatty acid oxidation) (Liesa and Shirihai, 2013), but also by regulating calcium stores (Duchen, 2000), intracellular signaling (Tait and Green, 2012) and apoptosis(Pradelli et al., 2010). Given the prominent role mitochondria play in nearly all facets of cellular functions, accurate determination of mitochondrial content is central in biomedical research. Many studies examining the changes in mitochondrial content under experimental or pathologic conditions limit the approach to utilizing Immunoblots with antibodies to two or more mitochondrial proteins as a surrogate marker for mitochondrial mass or oxidative phosphorylation (OXPHOS) complex activity (Choi et al., 2018; Matic et al., 2018; Nakashima-Kamimura et al., 2005; Nishimura et al., 2018; Velazquez-Villegas et al., 2018). Alternately, cell-permeable cationic dyes are often employed to measure the volume, relative membrane potential and superoxide status of mitochondria in primary or cultured cells via flow cytometry or fluorescence microscopy, but these dyes mis-localize under certain experimental conditions and need extensive calibration and proper controls in order to be informative (Padman et al., 2013; Puleston, 2015; Salvioli et al., 1997). Discovery or ‘shotgun’ proteomics approaches can provide a more comprehensive picture of mitochondrial content but have higher sample requirements in terms of processing to guarantee consistent detection of targeted, biologically selected mitochondrial proteins (e.g., mitochondrial-encoded vs nuclear-encoded electron transport chain components). While immunoblotting and other immunoassays target specific proteins of interest, quantitative immunoblotting approach is limited by the narrow linear relationship between the amount of the protein target and the intensity of the band as well as sensitivity to the target protein; in order to be valid, quantitative analyses must include a calibration curve and validation of specificity for each antibody used (Gilda et al., 2015; Murphy and Lamb, 2013). Further, the number of protein targets that can be analyzed on a immunoblot is limited, particularly without stripping and re-probing. A change in the amount of a single protein in a respiratory complex or in the tricarboxylic acid (TCA) cycle does not necessarily correspond to a change in the abundance or activity of the entire supercomplex; much less reflect the change in the mass or function of a complex organelle such as the mitochondria (Bezawork-Geleta et al., 2018; Fernandez-Vizarra et al., 2009; Guerrero-Castillo et al., 2017; Song et al., 2018). Finally, immunoblotting to quantify mitochondrial proteins is low throughput and the number of experimental samples that can be feasibly compared by immunoblot is limited. To address these methodological limitations and improve throughput and specificity of existing targeted mitochondrial quantification tools, we developed the mouse MitoPlex. MitoPlex is a targeted proteomics-based assay that utilizes scheduled Multiple Reaction Monitoring liquid chromatography mass spectrometry (MRM LC-MS) to quantify the total amount of a curated set of mouse mitochondrial proteins designed to reflect important components of mitochondrial function. The 37 proteins selected for the assay comprise both nuclear and mitochondrial-encoded components of the OXPHOS respiratory chain complexes, TCA cycle, mitochondrial protein transport, ROS reduction, mitochondrial DNA transcription, fatty acid oxidation, protein quality control and mitochondrial dynamics **(Table 1)** (Bezawork-Geleta et al., 2017; Diaz, 2010; Fernandez-Vizarra and Zeviani, 2015; Fiedorczuk and Sazanov, 2018; Jonckheere et al., 2012; Kang et al., 2018; Kerner and Hoppel, 2000; Nunes-Nesi et al., 2013; Pinti et al., 2016; Sebastian et al., 2017; Shokolenko and Alexeyev, 2017). The synergistic measurements of metabolites are helpful either via targeted assays, as we do in this study or in larger discovery metabolomic studies. Thus, to increase the power of the assay, we developed a reproducible protocol for isolating both metabolites and proteins from the same cell or tissue (in this case muscles, heart and liver) samples for parallel, focused mass spectrometry analysis, increasing the number of parameters that can be measured by our method and presenting a comprehensive reflection of metabolic status of the cells or tissues analyzed.

**Table 1.** MitoPlex Target Proteins

| Entry Name | Protein Name | Localization | Function |
| --- | --- | --- | --- |
| P08249\|MDHM | Malate dehydrogenase | matrix | TCA |
| P17710\|HXK1 | Hexokinase-1 | outer membrane | TCA |
| P35486\|ODPA | Pyruvate dehydrogenase E1 component subunit alpha | matrix | TCA |
| P47857\|PFKAM | ATP-dependent 6-phosphofructokinase | cytoplasm | TCA |
| P54071\|IDHP | Isocitrate dehydrogenase | matrix | TCA |
| P97807\|FUMH | Fumarate hydratase, mitochondrial | matrix | TCA |
| Q05920\|PYC M | Pyruvate carboxylase | matrix | TCA |
| Q60597\|ODO1 | 2-oxoglutarate dehydrogenase | matrix | TCA |
| Q99KI0\|ACON | Aconitate hydratase | matrix | TCA |
| Q9CZU6\|CISY | Citrate synthase | matrix | TCA |
| Q8K2B3\|SDHA | Succinate dehydrogenase flavoprotein subunit | inner membrane | TCA/OXPHOS |
| Q9CQA3\|SDHB | Succinate dehydrogenase iron-sulfur subunit | inner membrane | TCA/OXPHOS |
| Q9CZB0\|C560 | Succinate dehydrogenase cytochrome b560 subunit | inner membrane | TCA/OXPHOS |
| P00158\|CYB | Cytochrome b | inner membrane | OXPHOS |
| P00405\|COX2 | Cytochrome c oxidase subunit 2 | inner membrane | OXPHOS |
| P03921\|NU5M | NADH-ubiquinone oxidoreductase chain 5 | inner membrane | OXPHOS |
| P03930\|ATP8 | ATP synthase protein 8 | inner membrane | OXPHOS |
| P19783\|COX41 | Cytochrome c oxidase subunit 4 | inner membrane | OXPHOS |
| Q03265\|ATPA | ATP synthase subunit alpha | inner membrane | OXPHOS |
| Q8R1I1\|QCR9 | Cytochrome b-c1 complex subunit 9 | inner membrane | OXPHOS |
| Q91WD5\|NDUS2 | NADH dehydrogenase iron-sulfur protein 2 | inner membrane | OXPHOS |
| Q9CZ13\|QCR1 | Cytochrome b-c1 complex subunit 1 | inner membrane | OXPHOS |
| Q9D6J5\|NDUB8 | NADH dehydrogenase 1 beta subcomplex subunit 8 | inner membrane | OXPHOS |
| Q9DB77\|QCR2 | Cytochrome b-c1 complex subunit 2 | inner membrane | OXPHOS |
| Q9DCX2\|ATP5H | ATP synthase subunit d | inner membrane | OXPHOS |
| O35857\|TIM44 | Mitochondrial import inner membrane translocase TIM44 | inner membrane | Protein Transport |
| P09671\|SODM | Superoxide dismutase | matrix | ROS Reduction |
| Q8BK08\|TMM11 | Transmembrane protein 11 | inner membrane | Protein Transport |
| Q9CZW5\|TOM70 | Mitochondrial import receptor subunit TOM70 | outer membrane | Protein Transport |
| Q9D880\|TIM50 | Mitochondrial import inner membrane translocase TIM50 | inner membrane | Protein Transport |
| Q9DCC8\|TOM20 | Mitochondrial import receptor subunit TOM20 | outer membrane | Protein Transport |
| P40630\|TFAM | Transcription factor A | matrix | mtDNA Transcription |
| P52825\|CPT2 | Carnitine O-palmitoyltransferase 2 | inner membrane | Fatty Acid Transport |
| P58281\|OPA1 | Dynamin-like 120 kDa protein | inner membrane | Dynamics |
| Q811U4\|MFN1 | Mitofusin-1 | outer membrane | Dynamics |
| Q8CGK3\|LONM | Lon protease | matrix | Protein Quality Control |
| Q8K1M6\|DNM1L | Dynamin-1-like protein | outer membrane | Dynamics |

As proof of principle for the biological utility of MitoPlex, we analyzed two well-characterized mouse models: murine C2C12 myoblast cell line and the four major metabolically different C57BL/6J mouse tissues: heart, gastrocnemius muscle, soleus muscle and liver. C2C12 cells were originally isolated from the thigh muscle of a mouse, and in full-serum growth media rapidly proliferate and maintain an undifferentiated myogenic progenitor state; however, when exposed to low-serum differentiation media they undergo growth arrest and fuse into mature multi-nucleated myotubes (Burattini et al., 2004). Myoblasts are relatively quiescent cells that rely mainly on glycolysis. Upon differentiation, their mitochondria increase OXPHOS capacity by increasing the number of mitochondria per cell to handle the increased energy demand imposed by the contractile machinery (Sin et al., 2016), representing a robust system for demonstrating the utility of MitoPlex analysis *in vitro*. To test the ability of MitoPlex to report on changes in metabolic status under experimental conditions known to perturb mitochondrial function, we tested the effects of statins on C2C12 differentiation. Statins are a class of cholesterol-lowering drugs that inhibit 2-hydroxymethylglutaryl-coenzyme A (HMG-CoA) reductase that are widely prescribed to patients with cardiovascular disease, with a remarkably good safety profile. However, one of the more severe side effects is statin-mediated myopathy, caused primarily by mitochondrial dysfunction through the inability to generate Coenzyme Q_10_ (CoQ_10_)/Ubiquinone, resulting in reduced ATP production and depletion of mitochondria (Ramachandran and Wierzbicki, 2017). Statin myopathy most commonly presents as muscle pain, but in rare instances life-threatening rhabdomyolysis can develop.

Previous work has demonstrated that statin-induced myopathy is associated with mitochondrial myotoxicity and suppression of differentiation of C2C12 cells into myotubes (Baba et al., 2008; Schirris et al., 2015). Statins block the reduction of HMG-CoA to mevalonate, an essential precursor for the synthesis of cholesterol as well as for CoQ_10_. Addition of mevalonate has been shown to restore the ability of C2C12 cells to form tubes and implicitly, the ability to synthesize CoQ_10_ (Baba et al., 2008). A number of medical trials to supplement statin therapy with CoQ10 to relieve myalgia have been undertaken over the last several years, but the results are often contradictory and equivocal, possibly due to poor gastric uptake and delivery to tissues, as well as the poor diffusion of exogenous CoQ_10_ across plasma membranes and into mitochondria (Kaikkonen et al., 2002). In this study we use MitoPlex to interrogate whether CoQ supplementation restores myoblast differentiation that is blocked by when treated with statins (Banach et al., 2015; Qu et al., 2018).

To demonstrate the utility of our assay in analyzing tissue samples, we chose to focus on four tissues with discrete metabolic requirements. The heart and soleus (fast-twitch) skeletal muscle primarily utilize fatty acid oxidation as a source of fuel and generally maintain relatively high amounts of mitochondria per microgram of protein, whereas the gastrocnemius muscle (slow-twitch) is a more glycolytic fiber type and the liver is an organ that uses both glucose and fatty acids to support energy metabolism. Furthermore, all four of the tissues have differential energy requirements linked to their function and are therefore expected to exhibit different levels of mitochondrial proteins to reflect this activity (Grynberg and Demaison, 1996; Kummitha et al., 2014; Lee et al., 2017; Vitorino et al., 2007).

## Results

### Technical Performance of MitoPlex Assay

To establish the detection performance and reproducibility of the MRM-MS assay, we prepared total lysates, as well as cytoplasmic and mitochondria-enriched fractions from mouse C2C12 cells (N=3 replicate cultures) and soleus muscle homogenates (N=3 separate mice). Example peak groups for the quantification of the cytosolic glycolytic enzyme phosphofructokinase (PFK) and the mitochondrial tricarboxylic cycle enzyme aconitase (ACON) demonstrate clear enrichment of these proteins in the respective fractions of mouse soleus muscle samples (**Fig. 1A, B**). In all, 30 mitochondrial proteins were reliably quantified by the MRM-MS assay across the different fractions **(Fig. 1B)**, with the majority demonstrating excellent coefficient of variation (%CV) across technical replicates, particularly in the total lysate and mitochondrial preparations **(Fig. 1C, D)**. Poorer %CV in the cytosolic preparations likely reflects the expected experimentally induced reduction in mitochondrial protein content in this fraction and thus the volatility of quantification of MitoPlex proteins near their lower limit of detection / quantification levels.

**Figure 1.**
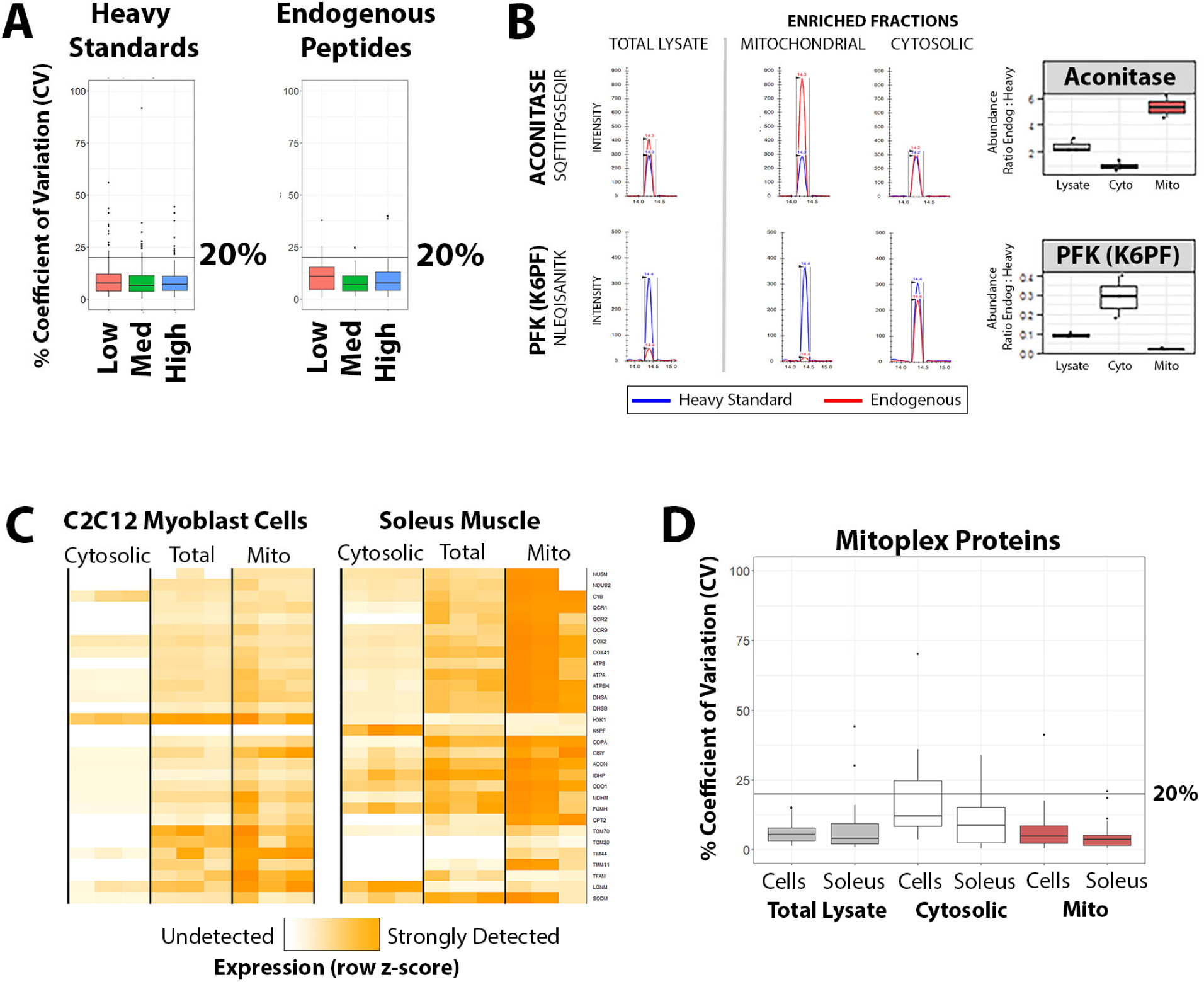
Technical Performance of MitoPlex Assay. **(A)** Heavy Stable Isotope Labelled reference peptides were spiked into C2C12 skeletal myoblast lysate at increasing concentrations relative (Low, 1:500 dilution; Medium 1:250 dilution; High 1:100 dilution). Stability of quantification was assessed using the coefficient of variation (standard deviation / mean * 100) across three technical replicates and assessed separately for the heavy standards (left panel) and the endogenous target peptides (right panel). **(B)** Example extracted ion chromatograms in soleus muscle samples for the heavy (blue) and light (red) peptides derived from the mitochondrial protein Aconitase and the cytosolic protein Phosphofructokinase (PFK/K6PF) are displayed in the hahahaleft panel for each of three sample preparation strategies (total lysate, crude mitochondrial enrichment, and crude cytosolic enrichment). Boxplots of the final protein quantification for these targets, calculated as the average ratio of endogenous to heavy observations of each fragment detected by the assay, are shown in the right panel. **(C)** Protein abundance heatmap for all MitoPlex proteins detected in C2C12 Myoblasts and Soleus Muscle cells within the difference sample preparations, Expression is displayed as row z-score, ranging from undetected / below lower limit of quantification (white) to highest in the dataset (dark orange). **(D)** Quantitative %CV for all proteins measured across the different sample preparation strategies, separated in to C2C12 Myoblasts (Cells) and Soleus Muscle (Soleus) groups.

### MitoPlex Characterization of Metabolic Differences in C2C12 Myotube, Myoblast and ρ^0^ Cells

To assess the ability of MitoPlex to report accurately on the changes in mitochondrial mass and content *in vitro*, we utilized a well-studied model of muscle development, myoblast cell line C2C12 (McMahon et al., 1994). When grown in differentiation media, these cells fuse and form functional myotubes that produce more ATP than the undifferentiated myoblasts; the response to this energy requirement is met by an increase in mitochondrial mass and OXPHOS capacity (Sin et al., 2016) **(Fig. 2A, B)**. In parallel, we established the ρ^0^ C2C12 cell line by depleting the myoblasts of mitochondrial DNA (mtDNA) by culturing the cells in a low concentration of ethidium bromide for 8 weeks, resulting in non-respiring mitochondria and a complete reliance on glycolysis for ATP production (Schubert et al., 2015) **(Fig. 2A, B).** The differences in mitochondrial content and activity in these cell lines were readily detected by the OXPHOS antibody cocktail (consisting of antibodies to NDUFB8, SDHB, UQCRC2, MTCO1 and ATP5A proteins) and reflected in Seahorse Bioanalyzer traces **(Fig. S1, 2A, B)**. In a MitoPlex comparison of myotubes and undifferentiated myoblasts, there was a clear increase in nearly all mitochondrial proteins per microgram of protein in the myotube whole lysate samples, with a notable exception of HXK1. This is consistent with a preference for oxidative phosphorylation of fatty acids in the differentiated cells as a fuel source **(Fig. 2C, D)**. Furthermore, in a comparison between myoblasts and ρ^0^ cells, we could not detect the peptides of mitochondrial DNA-encoded proteins NU5M, CYB, COX2 and ATP8 by MitoPlex in the ρ^0^ lysates, demonstrating that MitoPlex-generated data was accurately reproducing known results. Concomitantly, there was an increase in the mitochondrial proteins involved in the TCA cycle, reflecting the dysregulated metabolism in the OXPHOS-deficient ρ^0^ cells (**Fig. 2C, D**).

**Figure 2.**
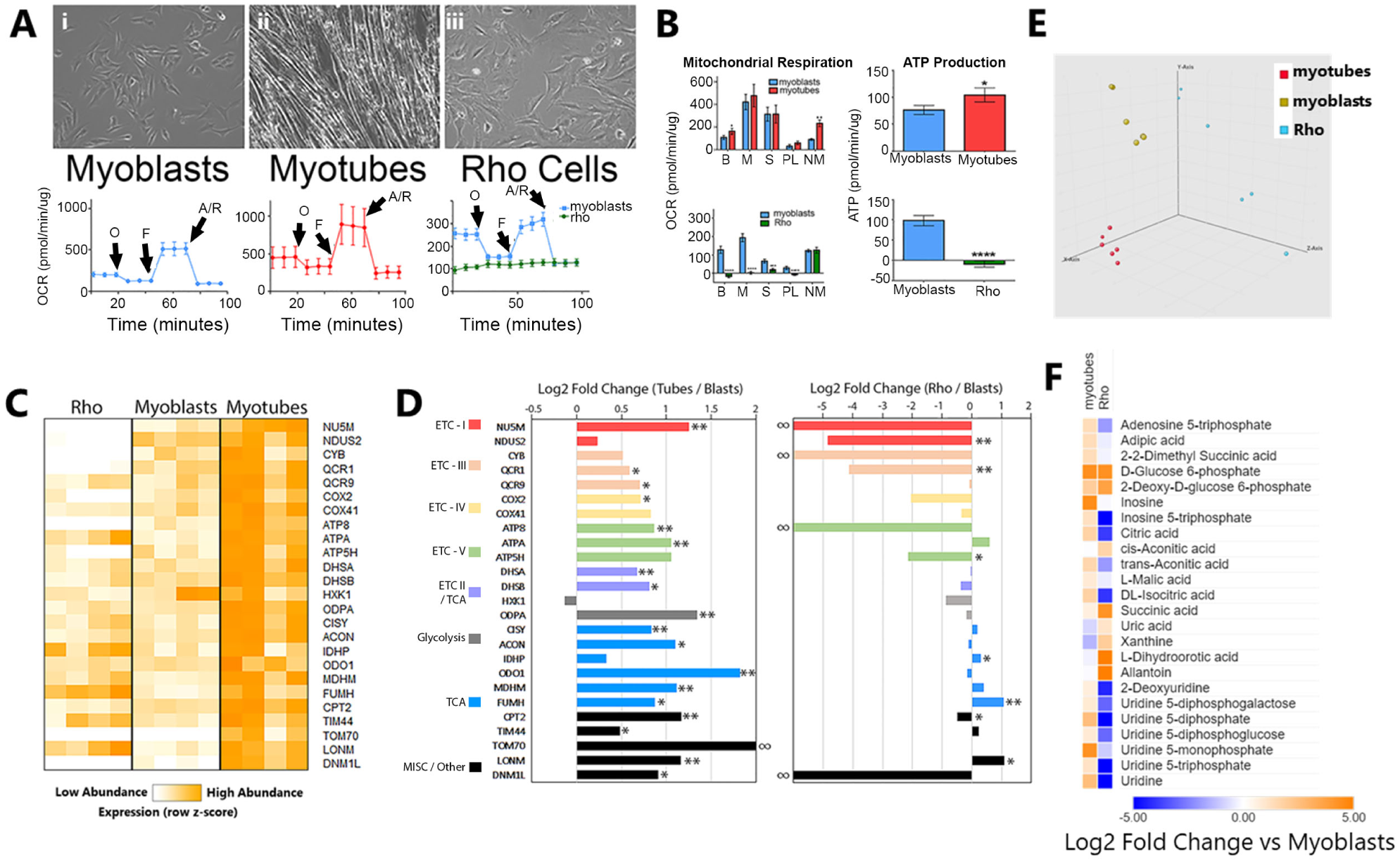
Characterization of C2C12 myoblasts, myotubes and Rho by MitoPlex. **(A)** Phase contrast microcopy images representative of C2C12 myoblasts (i), myotubes (ii) and (iii) Rho cells. **(B)** Respirometry trace and respiratory rates from C2C12 myoblast, myotube and Rho cells subjected to mitochondrial stress test; One-way analysis of variance (ANOVA) vs myoblasts, *p<0.05, **p<0.01, ***p<0.001, ****p<0.0001, values are means ±SD, n=3. B=basal respiration, M=maximal respiration, S=spare capacity, PL=proton leak, NM=non-mitochondrial respiration **(C)** Quantification of the results from **(B)**, values are means ±SD (n=4), unpaired t-test, *p<0.05 vs myoblast cells. **(C)** Protein abundance heatmap for all MitoPlex proteins detected in C2C12 myoblasts, myotubes and Rho cells **(D)** Comparison of Log2-fold changes in mitochondrial proteins between myotubes and myoblasts and Rho cells and myoblasts as reported by MitoPlex; values are means ±SD (n=4), unpaired t-test, *p<0.05 **p<0.01. **(E)** Principal component analysis of metabolites isolated from C2C12 myoblast, myotube and Rho cells (n=4). **(F)** Heat map analysis of LC-MS metabolomics comparing the myotubes vs myoblasts and Rho cells vs blasts. The row displays metabolite and the column represents the samples. Metabolites significantly decreased were displayed in blue, while metabolites significantly increased were displayed in red. The brightness of each color corresponded to the magnitude of the difference when compared with myoblasts.

Consistent with MitoPlex protein results, the metabolomic analysis of the three cell types revealed distinctly different profiles, segregating clearly by principal component analysis **(Fig. 2E)**. We detected statistically significant differences in 125 of the 218 metabolites assessed in the myotube and ρ^0^ cells as compared to myoblasts **(Fig. 2F)** supporting widespread metabolic differences between the different cell phenotypes.

### MitoPlex Characterizes the Block in Differentiation of Myoblasts in the Presence of Simvastatin and CoQ

We next examined the effect of Simvastatin with or without CoQ1 supplementation on myoblast differentiation and metabolic profile. Only the vehicle-treated cells were able to completely differentiate into myotubes based on visual inspection **(Fig. 3A)**. Concomitant with impaired tube formation, all three treatments (CoQ1, Simvastatin, Simvastatin+CoQ1) suppressed physiological measurements of basal respiration, spare respiratory capacity and maximal respiration **(Fig. 3B)**. Interestingly, despite clear functional differences in mitochondrial oxidation, the OXPHOS immunoblot only detected a change in SDHB (Complex II) protein levels between groups. **(Fig. 3C)**. In stark contrast, the MitoPlex assay detected decreased levels of nearly all members of the ETC, TCA, glycolysis and other mitochondrial proteins included in the assay when we treated cells with simvastatin, and this effect was dramatically amplified when CoQ_1_ was added **(Fig. 3D)**.

**Figure 3.**
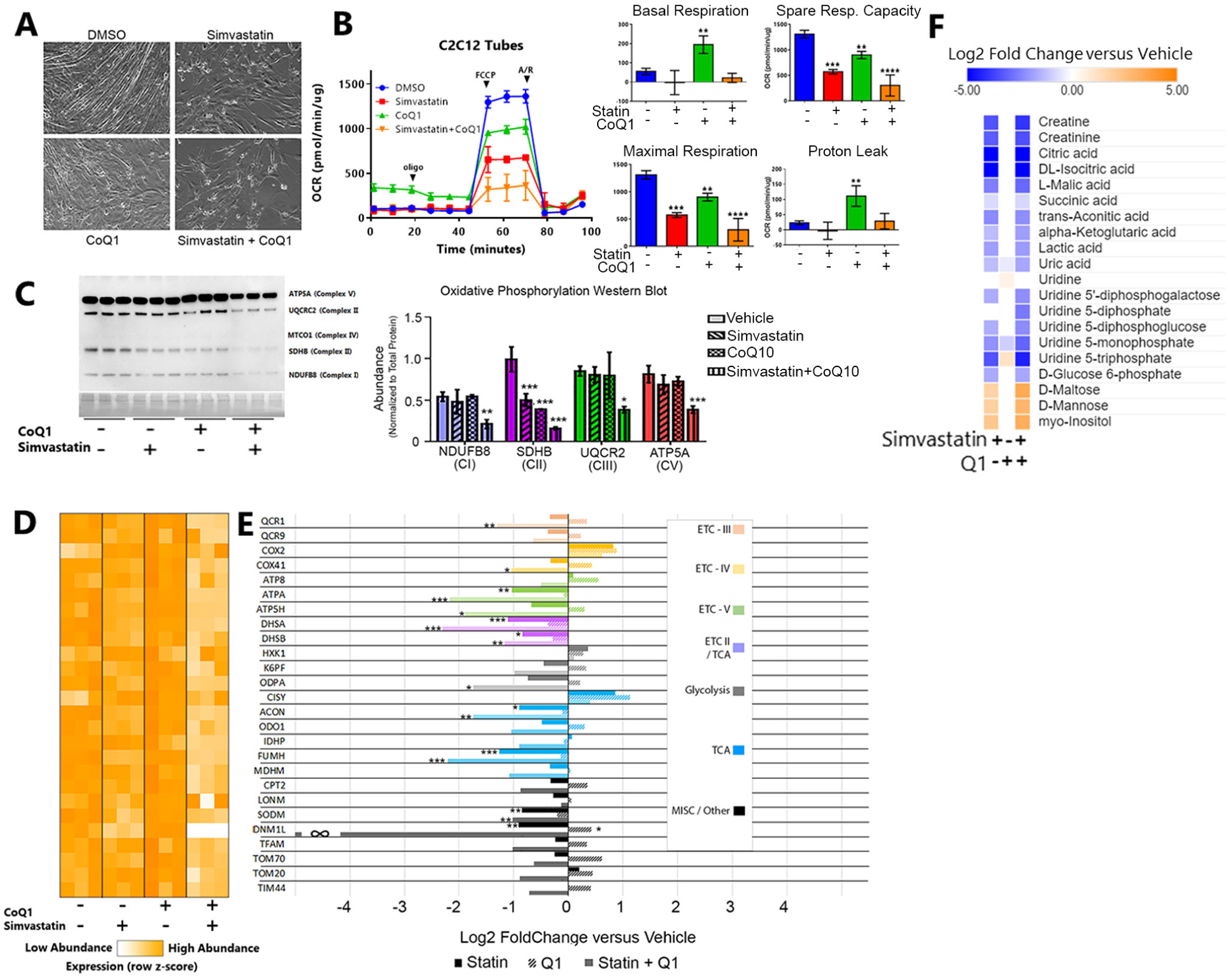
Comparison of Characterization of Effects of Simvastatin on C2C12 differentiation by Immunoblotting and MitoPlex. **(A)** Phase contrast microcopy images representative of C2C12 myotubes 6 days post differentiation in the presence of DMSO, 2uM simvastatin, 5uM CoQ1, or both. **(B)** Respirometry trace and respiratory rates from C2C12 myotubes differentiated in the presence DMSO, 2uM simvastatin, 5uM CoQ1, or both. One-way analysis of variance (ANOVA) vs DMSO, *p<0.05, **p<0.01, ***p<0.001, ****p<0.0001 are means ±SD, n=3. **(C)** Immunoblot analysis of whole cell lysates with the OXPHOS antibody cocktail and quantification of the results, normalized to Ponceau staining. One-way analysis of variance (ANOVA) vs DMSO, *p<0.05, **p<0.01, ***p<0.001, ****p<0.0001; representative immunoblot is shown, values are means ±SD, n=3). **(D)** Protein abundance heatmap for all MitoPlex proteins detected in C2C12 myotubes differentiated in the presence DMSO, 2uM simvastatin, 5uM CoQ1, or both. Expression is displayed as row z-score, ranging from undetected / below lower limit of quantification (white) to highest in the dataset (dark orange). **(E)** Comparison of Log2-fold changes in mitochondrial proteins between myotubes and myoblasts and Rho cells and myoblasts as reported by MitoPlex; values are means ±SD (n=3), unpaired t-test, *p<0.05 **p<0.01. **(F)** Heat map analysis of LC-MS metabolomics comparing

Metabolomic analysis of the C2C12 cells differentiated in the presence of simvastatin or simvastatin and CoQ1 as compared to the cells differentiated with DMSO further corroborated the inability of these cells to assemble high-functioning mitochondria, indicated by a decrease in the levels of creatine and creatinine, a decrease in intermediates of the TCA and glycolysis, (**Fig. 3E)**. Additionally, uric acid metabolism and nucleoside synthesis intermediates were decreased in the presence of simvastatin, while sugars like mannose and maltose were increased, as was myo-inositol, even as glycolytic intermediates decreased, indicating a dysregulated sugar metabolism. By contrast, addition of CoQ_1_ by itself did not drastically alter the metabolite profile as compared to DMSO treated cells, implying that the block in differentiation may occur independent of mitochondrial and metabolic remodeling.

### MitoPlex Accurately Differentiates Metabolic Differences between Tissues

MitoPlex performance was next evaluated in murine tissues, specifically tissue homogenates of mouse heart, gastrocnemius, liver and soleus skeletal muscle comparing the OXPHOS antibody cocktail versus MitoPlex. Immunoblot analysis suggests that the heart, liver and soleus muscle maintain a high level of mitochondrial protein per microgram of tissue, while the glycolytic gastrocnemius has less mitochondrial content. The blot also shows that the gastrocnemius maintains very low or undetectable levels of expression of Complex I and Complex III proteins at this protein load (20 ug/lane) **(Fig. 4A)**. MitoPlex assay confirms that the heart and soleus generally contain more mitochondrial protein per microgram than gastrocnemius or liver, but in contrast with the Immunoblot, MitoPlex clearly detects the presence of subunits of Complex I and Complex III **(Fig. 4B)**. Extending beyond the OXPHOS cocktail targets, the assay clearly reflects the fuel preferences of these tissues, as seen in the expression of KPF6, the enzyme responsible for the first step of glycolysis, which was present in much higher amount in the gastrocnemius muscle compared to the oxidative soleus muscle or the other tissues **(Fig. 4B, D)**; and CPT2, the enzyme responsible for delivering fatty acids to mitochondrial matrix for beta-oxidation, was present at a higher level in the liver, a major site of fatty-acid oxidation in the organism (Guillou et al., 2008; Webb et al., 2017) **(Fig. 4B, C)**. As expected, the comparison between the two different skeletal muscle types, the oxidative soleus fibers contained a higher level of mitochondrial proteins as compared to the gastrocnemius fibers **(Fig. 4D).** Interestingly, a pattern of tissue specific elevation of certain proteins (TFAM, SODM, and PYC) was observed, likely reflecting unique aspects of mitochondrial biology in these tissues.

**Figure 4.**
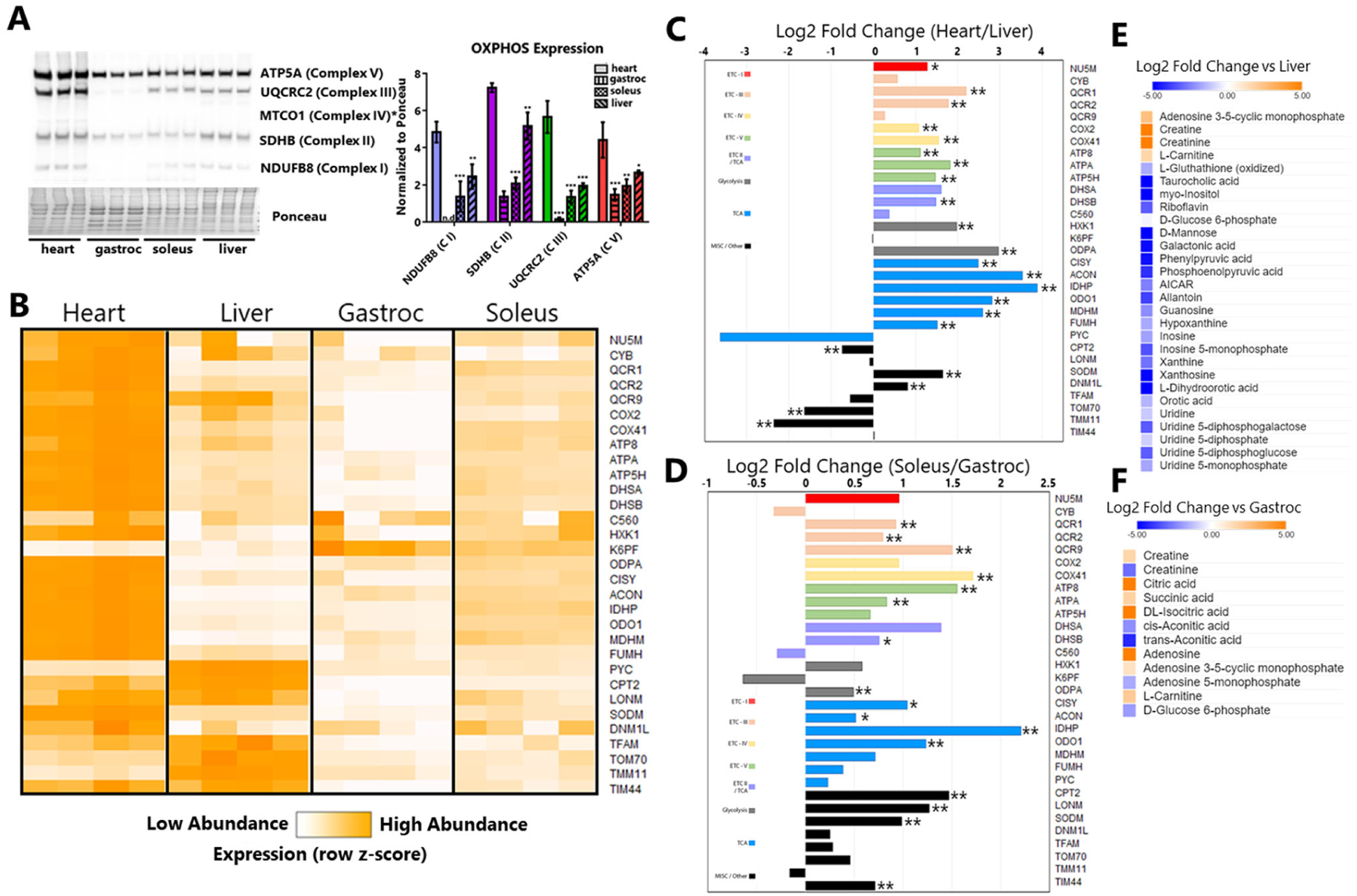
Characterization of murine heart, liver, gastrocnemius and soleus tissue by MitoPlex. **(A)** Immunoblot analysis of whole tissue lysates with the OXPHOS antibody cocktail and the quantification of the results, normalized to Ponceau staining. One-way analysis of variance (ANOVA) vs DMSO, *p<0.05, **p<0.01, ***p<0.001; representative immunoblot is shown, values are means ±SD, n=3). **(B)** Protein abundance heatmap for all MitoPlex proteins detected in tissues. Expression is displayed as row z-score, ranging from undetected / below lower limit of quantification (white) to highest in the dataset (dark orange). **(C)** Comparison of Log2-fold changes in mitochondrial proteins between heart and liver tissue as reported by MitoPlex; values are means ±SD (n=4), unpaired t-test, *p<0.05 **p<0.01. **(D)** Comparison of Log2-fold changes in mitochondrial proteins between soleus and gastrocnemius tissue as reported by MitoPlex; values are means ±SD (n=4), unpaired t-test, *p<0.05 **p<0.01. **(E)** Heat map analysis of LC-MS metabolomics comparing differences in the heart vs liver. **(F)** Heat map analysis of LC-MS metabolomics comparing differences in the soleus vs gastrocnemius.

Concomitantly extracted metabolomic analysis of the four tissues via the MitoPlex tissue processing protocol corroborated the MitoPlex results. There were 111 significantly different metabolites across the four different tissues, with multiple tissue-specific signatures consistent with the physiological function of the corresponding tissue **(Fig. 4E-F)**.

### MitoPlex Characterization of Differential Effects of Simvastatin on Organs

To demonstrate the ability of MitoPlex can quantify changes in an *in vivo* system, we investigated the effect of 10 days of simvastatin treatment on cardiac muscle, liver, gastrocnemius and soleus muscle mitochondrial content (n=4). Interestingly, MitoPlex revealed a trend toward a disparate response to simvastatin treatment in the heart and liver as compared to the skeletal muscles (**Fig. 5**). In liver and heart, nearly all mitochondrial proteins displayed a positive trend compared to vehicle controls (**Fig. 5A, B)**. By contrast, the mitochondrial proteins of gastrocnemius and soleus muscles displayed a largely negative trend (**Fig. 5A, B)**. The metabolite data further corroborated these trends, and the major metabolites of the TCA cycle correlated to the protein members in terms of relative amounts; notably, the metabolites tended to increase in the hearts and livers of statin treated mice and decrease in the gastrocnemius and soleus muscles (**Fig. 6)**.

**Figure 5.**
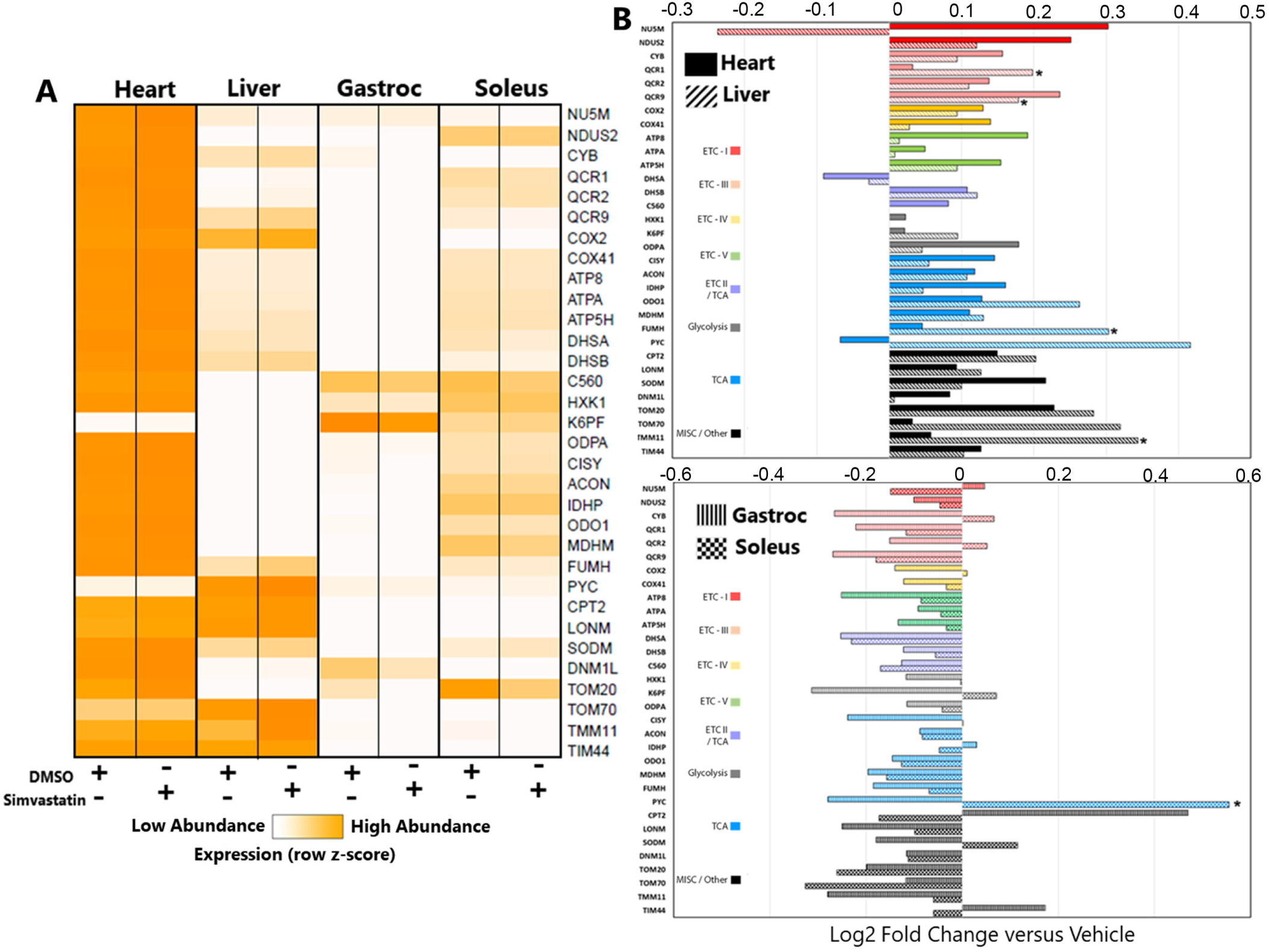
MitoPlex Characterization of Differential Effects of Simvastatin on Organs. **(A)** Protein abundance heatmap for all MitoPlex proteins detected in organs of mice treated for 10 days with either DMSO or 20 mg/kg simvastatin. Expression is displayed as row z-score, ranging from undetected / below lower limit of quantification (white) to highest in the dataset (dark orange). **(B)** Log2-fold changes in mitochondrial proteins in the tissues of simvastatin-treated mice (vs. vehicle) as reported by MitoPlex; values are means ±SD (n=4), unpaired t-test, *p<0.05. **(C)** Log2-fold changes in mitochondrial proteins and metabolites of the TCA cycle in the tissues of mice treated with simvastatin (vs. vehicle). Proteins are labeled in red and metabolites in blue.

**Figure 6.**
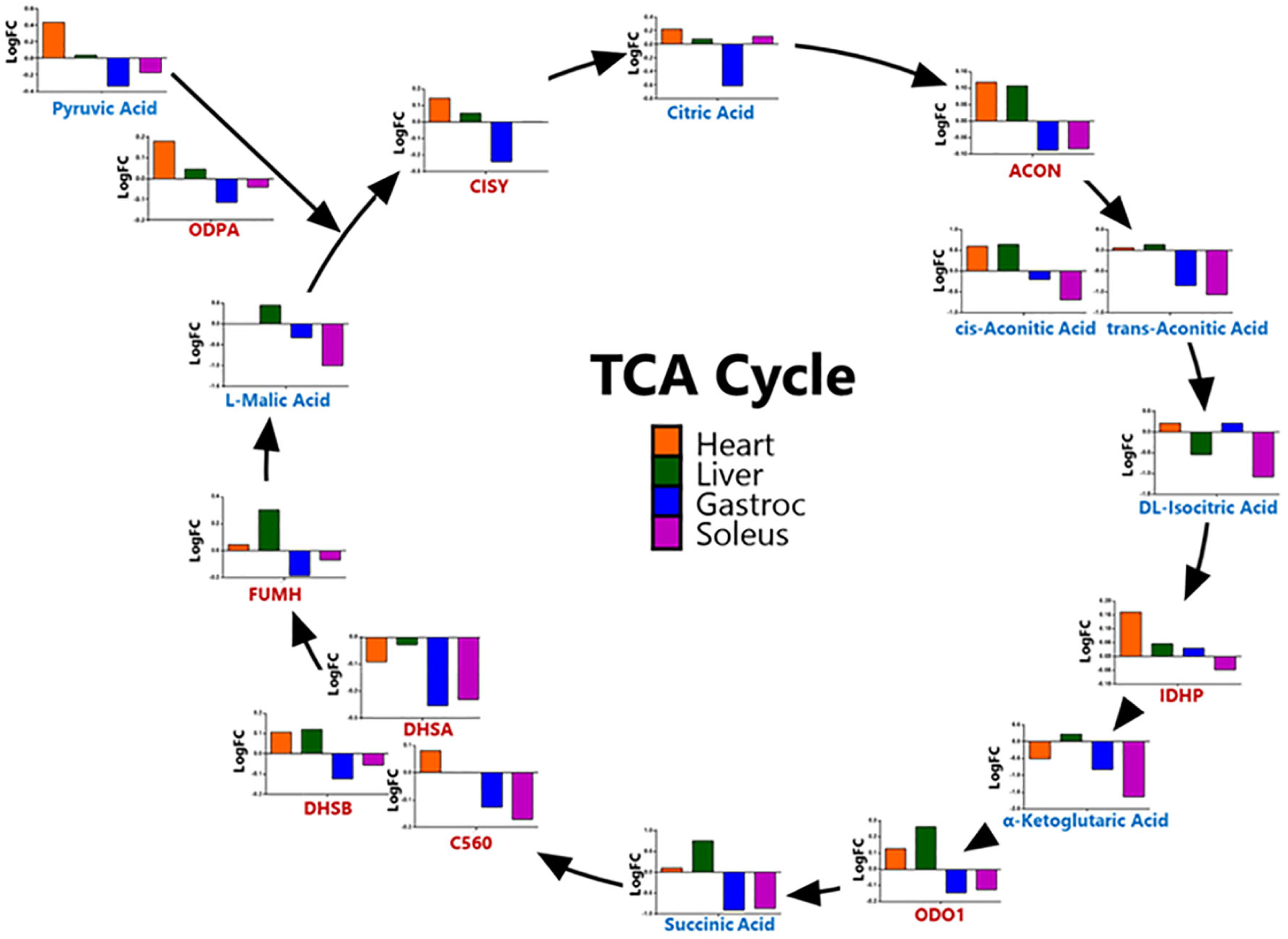
MitoPlex Characterization of Effects of Simvastatin on TCA Cycle Enzymes and Metabolites. Log2-fold changes in mitochondrial proteins and metabolites of the TCA cycle in the tissues of mice treated with simvastatin (vs. vehicle). Proteins are labeled in red and metabolites in blue.

## Discussion

Here we report a novel mass spectrometry assay for quantifying mitochondrial mass and metabolites *in vitro* and *in vivo.* The assay is highly sensitive, reproducible and far superior to the commonly used methods in profiling mitochondrial content. The modular nature of the assay allows for expansion and inclusion of other proteins of interest beyond those included in this iteration, and the ability to process the same sample for proteomic and metabolomic analysis allows for a comprehensive interrogation of the metabolic status of the cells or tissues analyzed. Our analysis of technical performance of the MitoPlex assay in mitochondria-enriched and total lysate samples demonstrated high precision across technical replicate preparations (CVs well below 20%), and also appropriate biological information by demonstrating depletion of mitochondrial proteins from cytosolic fractions.

We demonstrate the ability of MitoPlex to accurately reflect the changes in mitochondrial content *in* vitro, in the comparison between the myoblasts, myotubes and ρ^0^ cells as well as in the comparison of C2C12 cells treated with simvastatin and/or CoQ. The assay reflected differences in mitochondrial mass and metabolite content between cells at different differentiation stages, mitochondrial genome statuses (ρ^0^), and under different treatment conditions, clearly demonstrating the drastic increases in the protein and metabolite quantities when myoblasts form into functional myotubes, and effect that is blocked by statin and/or CoQ treatment. Importantly, immunoblot methods failed to detect similar decreases most mitochondrial proteins relative to the MitoPlex observations. Interestingly, the assay also indicated that there was an increase in Lon protease (LONM) in ρ^0^ cells, a previously unpublished finding (**Fig. 2C, D**). LONM is a mitochondrial protease responsible for clearing misfolded proteins and alleviating proteotoxic stress in the organelle; it is important for the mitochondrial unfolded protein response and through its degradation of PINK1, for suppressing mitophagy (Jin and Youle, 2013; Zurita Rendon and Shoubridge, 2018).

The ability to extract both proteins and metabolites from the same sample allows for a deeper analysis of the metabolic status of tissues or cells. For instance, in the C2C12 differentiation program, consistent with corresponding seahorse physiological assessment of mitochondrial function, the more metabolically active myotubes had higher whereas the ρ^0^ cells had less adenosine triphosphate (ATP) compared to the myoblasts. Consistent with higher reliance on glycolysis, ρ^0^ cells also had a higher level of 2-deoxy-D-glucose-6-phosphateand myoinositol, which stimulates uptake of glucose from the media (Dang et al., 2010). The loss of the respiratory electron transfer chain in ρ^0^ cells results in inhibition of dihydroorotate dehydrogenase (DHODH) and an inability to generate uridine and other pyrimidines. This was clearly reflected in the metabolite analysis of the ρ^0^ cells by the absence of uridine as well as its downstream metabolites, an increase in the level of the pyrimidine precursor L-dihydroorotic acid, and a decrease in cytidine and inosine triphosphate, reflecting impaired pyrimidine biosynthesis (Huang and Graves, 2003). Additionally, ρ^0^ cells displayed higher levels of xanthine, uric acid and allantoin, which are breakdown products of purines, likely due to the imbalance of purine and pyrimidine biosynthesis. By comparison, relative to myoblasts, myotubes displayed an overall increase in all metabolites, reflecting the higher metabolic rates in the myotubes, with a notable decrease in myoinositol levels, reflecting the preference for fatty acids to support increased ATP generation in myotubes (Shintaku et al., 2016). Interestingly, myotubes had higher levels of tryptophan and L-kynurenine and a lower level of 5-hydroxy-3-indoleacetic acid, indicating an increase in nicotinamide adenine dinucleotide (NAD+) synthesis as opposed to serotonin synthesis in tryptophan downstream metabolism (Cervenka et al., 2017).

Metabolite analysis of the differentiating C2C12 cells in the presence of simvastatin revealed that statistically significant decreases in metabolites such as succinic acid, uric acid, glucose and fructose, and xanthine that are consistent with the response of patients undergoing simvastatin therapy that are good responders (Trupp et al., 2012). Unfortunately, these same pathways appear important for the normal function of differentiated myotubes and thus, this effect may underlie the detrimental effect of statins on muscle function. Our data also suggest that adding CoQ to simvastatin-treated myoblasts does not rescue them but in fact is detrimental to cell survival under the stress conditions of differentiation. ROS serve as an important activator of the master mitochondrial biogenesis regulator PGC-1α (Yoboue and Devin, 2012), and when the antioxidant CoQ is added, mitochondrial biogenesis necessary to increase mitochondrial mass in differentiating cells may be blocked.

We also demonstrated the ability of the MitoPlex and metabolite workflow to detect phenotypic and treatment-induced differences in mitochondrial potential of various tissues. Comparison of heart, liver, soleus and gastrocnemius tissues recapitulated several known metabolic differences between these tissues. Of note are the relatively high levels of mitochondrial import machinery and the mitochondrial transcription factor TFAM present in the liver as compared to the other 3 tissues, reflecting the high rate of mitochondrial protein turnover in the liver (Chan et al., 2015). Compared to the other tissues, the heart contains a high level of SODM, essential for managing oxidative stress in the mitochondria, and likely a reflection of the high rate of oxidative phosphorylation in that organ (Younus, 2018). The liver also contained strikingly high levels of pyruvate carboxylase (PYC) **(Fig. 3B, C)**, reflecting the major role the organ plays in gluconeogenesis and lipogenesis (Jitrapakdee et al., 1996). In the comparison of the metabolite expression levels between heart and liver, clear functional differences were detected, as seen in the high levels of cAMP, creatine, creatinine and L-carnitine in the heart. These elevated metabolite levels are consistent with the fact that cAMP mediates the catecholaminergic control on heart rate and contractility; creatine and its degradation product creatinine are pivotal parts of the mitochondrial ATP shuttle; and L-carnitine is conjugated to fatty acids to enable their import into the mitochondria for beta-oxidation in cardiomyocytes (Wang et al., 2018; Zaccolo, 2009; Zervou et al., 2016). In contrast, and consistent with its distinct functions, the liver contained higher levels of taurocholic acid (liver bile salt), phenylpyruvic acid (phenylalanine metabolism restricted to liver and kidneys), phosphoenolpyruvic acid (gluconeogenesis), and riboflavin (vitamin storage) (Baker et al., 1964; Garibotto et al., 2002; Liu et al., 2018; Yang et al., 2009). Liver samples also contained higher levels of mannose, glucose-6-phosphate and galactonic acid, reflecting the differences in fuel utilization preferences between the heart (mostly fatty acids) and the liver (mix of glycolysis and fatty acid oxidation) (Rui, 2014; Stanley and Chandler, 2002; Wood et al., 1961). The liver is a major site of uric acid metabolism, and this is reflected by the higher amounts of intermediate metabolites in the sample, as compared to the heart (Maiuolo et al., 2016). Finally, the liver had much higher rates of uridine metabolic intermediates, as expected from the tissue where uridine is scavenged from the serum as well as synthesized *de novo* (Le et al., 2013). The comparison between the soleus and gastrocnemius muscles accurately reflected the differences between the oxidative and glycolytic fibers by the presence of higher amounts of TCA intermediates (apart from aconitase), L-carnitine and creatine in the soleus, and lower amounts of glucose-6-phosphate.

The high degree of precision of the assay allows for the detection of subtle changes in mitochondrial content and metabolite level in tissues; although the effect was small and did not achieve statistical significance for most analytes (3-9% average with 6-11%CVs), there was a clear and consistent trend between both proteins and metabolites indicating divergent response of different tissues to statin treatment. Specifically, while the heart and liver demonstrated nearly universal, albeit small, increases in protein and metabolite levels the skeletal gastrocnemius and soleus muscles exhibited consistent reductions in these same analytes. This differential response of cardiac and skeletal muscles to statins is consistent with previous reports, demonstrating that in human patients, cardiac atrial muscle oxidative capacity was enhanced relative to untreated counterparts while in deltoid skeletal muscle biopsies from patients experiencing statin-induced myopathy, oxidative capacity was diminished relative to untreated counterparts. The authors go on to link this differential effect of statin in these two tissue types to a potential discrepancy in redox buffering capacity and subsequent PGC1a activation (Bouitbir et al., 2012). Unfortunately, deltoid biopsies of patients treated with statin but not experiencing myopathy were not examined, and atrial samples and deltoid samples were not collected from the same patients, so intra-patient differences cannot be verified by this previous study. It remains to be seen whether, in humans, those prone to myopathy demonstrate similar or divergent response to statin between atrial and skeletal muscle tissues. Our data in mice indicate that, indeed, at least in otherwise normal mice treated for a relatively short duration there are clear trends toward divergence. The lack of statistical significance may be overcome by additional replicates, however it may also be important to consider that just as not all humans experience myopathy side effects with statins, mouse models of statin-induced myopathy to more severe or clinically relevant levels may require additional predisposing factors that render a mouse or human vulnerable to these detrimental side effects. Importantly, the synergy of the MitoPlex plus metabolites approach enables screening for factors that promote or exacerbate statin-mediated mitochondrial impairment in skeletal muscle models and may ultimately aid in identifying targets for adjuvant therapies that mitigate detrimental skeletal muscle side effects of this otherwise protective and powerful class of cardiovascular therapeutic drugs.

## Conclusion

The combination of MitoPlex protein quantification and targeted metabolite analysis primarily of the central carbon chain metabolism which drives ATP production, provides a detailed portrait of mitochondrial metabolism. This precise and reproducible assay requires a small amount of material, suitable for analysis of all mouse cells and tissues, and is a high throughput, focused, and powerful addition and complement to the current techniques in use for the study of mitochondrial biology in mouse models.

## LEAD CONTACT AND MATERIALS AVAILABILITY

Further information and requests for resources and reagents should be directed to and will be fulfilled by the Lead Contact, Aleksandr Stotland

## EXPERIMENTAL MODEL AND SUBJECT DETAILS

### Experimental Animals

Sixteen-week old male C57/BL6J were purchased from Jackson Laboratory (Cat#000664). All aspects of the experiments performed, including procurement, conditioning, quarantine, housing, management, veterinary care and disposal of carcasses follow the guidelines set down in the NIH Guide for the Care and Use of Laboratory Animals, and approved by the Cedars-Sinai Institutional Animal Care and Use Committee. All mice were housed in a temperature-controlled environment (23°C) with a 12-h light and 12-h dark (18.00–06.00) photoperiod and cared in facilities operated by the Department of Comparative Medicine at Cedars-Sinai Medical Center. The department is fully accredited by the Association for Assessment and Accreditation of Laboratory Animal Care (AAALAC) and provides a full range of clinical veterinary services including disease diagnosis, prevention, and treatment on a 24-hr basis for animal research. Animals were provided PicoLab High Energy Mouse Diet (PicoLab, Cat#5LJ5) and water *ad libitum*.

### C2C12 Cell Culture

C2C12 mouse skeletal myoblasts (ATCC, CRL-1772) were maintained in growth media consisting of DMEM (Gibco, 11995–073) containing 10% fetal bovine serum (Life Technologies, 16010–159) and antibiotic/antimycotic (Life Technologies, 15240–062). Cells were differentiated by allowing them to reach 100% confluence and switching them to differentiation medium consisting of DMEM containing 2% horse serum (Life Technologies, 16050–122) and antibiotic/antimycotic. After 6 days in differentiation media, cells appeared mostly as phase-bright fused myotubes. C2C12 ρ0 cells were generated by growing C2C12 myoblasts in growth media supplemented with 50 ng/ml of ethidium bromide (Sigma-Aldrich, E1510-10ML) and 50 µg/ml of uridine for 8 weeks, and the loss of the mitochondrial genome was confirmed by PCR. To characterize the changes in mitochondrial content in differentiating C2C12 cells treated with statin and to determine whether CoQ10 supplementation is beneficial under such treatment, we grew the myoblasts in differentiation media for 7 days in the presence of the vehicle DMSO, 2 µm simvastatin (Toronto Research Chemicals, S485000), 5 µm of a CoQ10 soluble analog (CoQ1, Sigma-Aldrich, C7956), or combined treatment. Doses and protocol were derived from previously published work (Chan et al., 2004).

## METHOD DETAILS

### Discovery Proteomic Analysis

To identify MS-detectable, informative peptides for development into the targeted MitoPlex assay, mitochondria were isolated from C2C12 cells by scraping into mitochondrial isolation buffer containing sucrose (250 mM; Sigma-Aldrich, 179949), EDTA (1 mM), HEPES (10 mM; Sigma-Aldrich, H3375, pH 7.4), and protease inhibitors (Roche, Cat#11697498001). Cells were then disrupted by nitrogen cavitation (Gottlieb and Adachi, 2000), and cell slurries were centrifuged at 950 x g for 5 min to pellet the nuclei and unbroken cells. Supernatants were collected and centrifuged again at 7000 x g for 15 min to pellet heavy membranes (mitochondria-enriched) fractions. Enriched mitochondrial pellets were lysed in 8M urea/50mM Tris-HCl, pH 8.0 with sonication (10 seconds at 70%power followed by 10 seconds on ice with a Thermo Fisher Scientific Qsonica Sonicator Q125) and BCA assay was performed to quantify the protein concentration (Thermo Fisher Scientific, Cat#23225). An aliquot of 50 µg of total protein was made and diluted to a final concentration of 2M urea using sequential incubations with 10mM dithiothreitol (DTT, Thermo Fisher Scientific, Cat#BP172-5) for 20 minutes at 37oC and 100mM iodoacetamide (IAA, Sigma-Aldrich, Cat#16125-25G) for 20 minutes at room temperature in the dark to dilute the samples while also reducing and alkylating cysteine residues. Trypsin/LysC (Promega, Cat#V5072) was added at a ratio of 1:40 total protein and samples were digested overnight. Subsequent peptides were desalted on NEST C18 tips (Thermo Fisher Scientific, Cat#NC0484000) and dried. To identify optimally detectable and quantifiable proteins, both data dependent and data independent acquisitions were performed on a Triple TOF 6600 Mass Spectrometer as described in supplemental methods. The raw data dependent acquisition file was searched using the transproteomic pipeline as described previously (Parker et al., 2016). The InterProphetParser file was imported into Skyline software for generation of a mitochondrial protein spectral library. All proteins identified with an iprophet probability >0.95 were manually inspected, and representative proteins from curated functional categories were selected for inclusion in the downstream targeted assay. Candidate peptides from each of these proteins were selected by interrogating fragment peak groups in Skyline, and at least two peptides for each target protein were selected based on overall intensity and signal-to-noise characteristics from the Skyline analysis. Stable isotope standard (SIS) labeled versions of these peptides were synthesized with heavy 13C 15N-labelled versions of their C-terminal amino acid (New England Peptide, Boston MA). Synthetic heavy peptides were pooled, and a preliminary series of unscheduled, targeted acquisition methods were exported from Skyline using predicted acquisition settings for a QTRAP 6500 mass spectrometer (Sciex) (Fu et al., 2018). Data were acquired as follows: Peptides were separated on a Prominence UFLCXR HPLC system (Shimadzu, Japan) with a Waters Xbridge BEH30 C18 2.1mm x 100mm, 3.5µm column (Waters) flowing at 0.25 mL/min and 36 °C coupled to a QTRAP® 6500 (SCIEX, Framingham, MA). Mobile phase A consisted of 2% ACN, 98% water, and 0.1% formic acid and mobile phase B of 95% ACN, 5% water, and 0.1% formic acid. After loading, the column was equilibrated with 5% B for 5 minutes. Peptides were then eluted over 30 minutes with a linear 5% to 35% gradient of buffer B. The column was washed with 98% B for 10 minutes and then returned to 5% B for 5 minutes before loading the next sample. Raw data were imported into Skyline in order to examine the quality of each synthetic peptide. Poorly detectable synthetic peptides (e.g., low signal to noise, low intensity) acquired with Skyline predicted parameters were further optimized for collision energy, de-clustering potential by direct injection onto the QTRAP 6500. Unscheduled acquisition was repeated following optimization to ascertain retention time.

### Scheduled Multiple Reaction Monitoring Liquid Chromatography Mass Spectrometry

Retention times were determined from the unscheduled acquisitions and used to generate a scheduled (+/− 2 minutes of expected RT), optimized acquisition list including both the labelled (heavy) and unlabeled (endogenous) versions of each peptide. For assessing initial performance, triplicate preparations of cytosolic, mitochondria-enriched, and total homogenate from both mouse soleus muscle and C2C12 myotube lysates were digested with trypsin, desalted, and spiked with heavy SIS peptides. For treatment and experimental studies, whole cell lysates and tissue homogenates were digested with trypsin, desalted, and combined with SIS peptides. All samples were analyzed with the final scheduled acquisition method on the QTRAP6500 as described above to assess quantitative sensitivity and precision of the assay.

### MRM Data Analysis

Raw MS-data were imported into Skyline and peptide peaks were manually analyzed for consistent retention time and co-elution of heavy and endogenous peptide transitions. Data were then exported, and a custom R script was generated to calculate coefficient of variation across all heavy transitions and to filter all fragments to include only those with a heavy peptide CV <20%. Specifically, the standard deviation of all observations of a heavy fragment across all samples was divided by its average, and fragments with >0.2 were removed from further analysis. Ratio of endogenous to heavy area ratios, as calculated from Skyline software, were averaged for all high quality fragments from a given peptide and subsequently the high quality peptides from a given protein to produce a final peak area ratio for each protein reliably quantified from a given set of experimental preparations.

### Metabolomic Analyses (dMRM)

Cell metabolite extractions were analyzed with an Agilent 6470A Triple quadrupole mass spectrometer, operating in negative mode, connected to an Agilent 1290 Ultra High-Performance Liquid Chromatography (UHPLC) system (Agilent Technologies, Santa Clara CA). Mobile phases consisted of HPLC or LCMS grade reagents. Buffer A is water with 3% methanol, 10mM tributylamine (TBA), and 15mM acetic acid. Buffers B and D are isopropanol and acetonitrile, respectively. Finally, Buffer C is methanol with 10mM TBA and 15mM acetic acid. The analytical column used was an Agilent ZORBAX RRHD Extend-C18 1.8µm 2.1 x 150 mm coupled with a ZORBAX Extend Fast Guard column for UHPLC Extend-C18, 1.8 µm, 2.1 mm x 5mm. The MassHunter Metabolomics dMRM Database and Method was used to scan for up to 219 polar metabolites within each sample (Agilent Technologies, Santa Clara CA). Resulting chromatograms were visualized in Agilent MassHunter Quantitative Analysis for QQQ. The final peaks were manually checked for consistent and proper integration. Statistical analysis was completed in Agilent Mass Profiler Professional (Agilent Technologies, Santa Clara CA). Samples were normalized to the single copy nuclear gene ApoB as detected by quantitative PCR (detailed protocol in the supplement) (Machado et al., 2015).

### Immunobloting

Whole cell lysates were obtained by applying RIPA buffer containing Tris pH 8.0 (50 mM; Sigma-Aldrich, T1503), NaCl (150 mM; Sigma-Aldrich, S7653), ethylene glycol tetraacetic acid [EGTA] (2 mM; Sigma-Aldrich, E4378), ethylenediaminetetraacetic acid [EDTA] (1 mM; Sigma-Aldrich, E4884), NP-40 (1%; Sigma-Aldrich, I3021), sodium deoxycholate (0.5%; Sigma-Aldrich, D6750), sodium dodecyl sulfate [SDS] (0.1%; Bio-Rad Laboratories Inc. Laboratories Inc., 161–0302) directly to adherent cells and scraping. Tissue samples were subjected to polytron homogenization in RIPA buffer at 4°C. Proteins were quantified using bicinchoninic acid assay (Sigma-Aldrich, B9643). Equal amounts of protein were run in 4–20% Tris-glycine SDS-PAGE gels (Life Technologies, Cat#EC6025) and transferred to nitrocellulose membranes. Membranes were blocked in 5% nonfat dry milk in Tris-buffered saline [TBS] containing Tris (20 mM) and NaCl (150 mM) pH-adjusted to 7.6 with 0.1% Tween-20 (Sigma-Aldrich, P1379) [TBS-T] for 1 hr at room temperature and then incubated in primary antibody diluted in 5% nonfat dry milk overnight at 4°C. OXPHOS complex antibody cocktail (1:250, Abcam, Cat#ab110413) was used for the detection of the respiratory chain proteins. Membranes were washed in TBS-T and incubated in horseradish peroxidase-conjugated anti-mouse (1:3000, KPL, Cat#074–1806) secondary antibody for 1 hr at room temperature. Membranes were washed again in TBS-T and developed with Clarity Western ECL Substrate (Bio-Rad Laboratories Inc. Laboratories Inc., Cat#170–5061) and imaged using a Bio-Rad Laboratories Inc. ChemiDoc XRS (Bio-Rad Laboratories Inc. Laboratories Inc.; Hercules, CA, USA). Densitometry was performed using ImageJ software (NIH) within the linear range of the antibodies.

### Respirometry

Intact cellular respirometry was conducted using Seahorse XFe24 extracellular flux analyzer. C2C12 myoblasts or ρ0 cells were seeded onto XF24 V7 cell culture plates or differentiated into myotubes for 7 days prior to the analysis. The cells were refreshed with bicarbonate-free DMEM containing 25 mM glucose, 1 mM sodium pyruvate, 2 mM glutamine and equilibrated for 1 hr at 37°C in a non-CO2 incubator. Oxygen consumption was subsequently monitored following sequential injection of oligomycin (1 µM), FCCP (1.25 µM) & antimycin/rotenone (1 µM/1 µM) (Seahorse XF Cell Mito Stress Test Kit, Agilent, Cat#103015-100). For normalizing respiration rates, cells were subsequently lysed in lysis buffer and protein concentration was determined using the bicinchoninic acid assay. Results were analyzed with the Seahorse Wave Desktop Software (Agilent).

### Mouse Simvastatin Treatment

Eight sixteen-week old C57/BL6 mice were injected with either vehicle (DMSO) or 20mg/kg of simvastatin (Toronto Research Chemicals, Cat#S485000) once a day intraperitoneally for 10 days. Organs were harvested and processed as described in Single Sample Preparation protocol.

### Single Sample Preparation Protocol for Cells

Adherent cells were seeded in 6-well culture plates for experimental treatments. Growth media was removed, and the cells were rinsed twice with 2ml of phosphate-free saline (Vetivex, Dechra NDC# 17033-492-01). Saline was removed and 800ul of metabolite extraction buffer was added to the cells (40% acetonitrile (Fisher Scientific, A996-1), 40% methanol (Fisher Scientific, A454-1) and 20% water (Fisher Scientific, W6500)). Plates were gently swirled on ice at 4C for 2 mins and supernatant was transferred into 1.5ml Eppendorf tubes. Supernatant was spun in a microcentrifuge (Thermo Scientific, Sorvall Legend Micro 21R) at 21,000g for 10 min at 4C to pellet cell debris and transferred to a new Eppendorf tube. Metabolites were precipitated by a 6 hr spin in SpeedVac concentrator (Thermo Scientific, Savant SPD2010). The cells were then rinsed with 4ml of saline, 2ml of saline with 10mg/ml bovine serum albumin (Sigma-Aldrich, Cat#A0281-10G) and lifted by scraping the cells with a cell scraper. Cells were pellet by spinning at 500g for 5 min at 4C, supernatant removed, the cells gently washed with BSA-free saline and pelleted once again. Following the spin, the cells were lysed in 200ul 8M urea/50mM Tris-HCl, pH 8.0 and processed for proteomic analysis as described above.

### Single Sample Preparation Protocol for Tissues

Mice were anesthetized by intraperitoneal injection of 5 mg of Fatal-Plus solution (sodium pentobarbital) (Vortech Pharmaceuticals, Ltd). After confirming the mouse was anesthetized by the lack of toe pinch and tail pinch response, the chest cavity was exposed, and the liver was pulled back to access the portal vein. The vein was cut to aid in exsanguination and the heart was flushed by injecting 10ml of saline into the apex of the heart with a 27-gauge syringe, at a steady rate for no longer than 1 minute. The heart was extracted and immediately placed in 800 ul metabolite extraction buffer and homogenized on ice with a 5 second pulse at 50% power using a pre-cooled Polytron homogenizer (PowerGen 125, Thermo Fisher Scientific, Waltham, MA). The homogenization process was repeated for the liver and skeletal muscle samples. The homogenized samples were then centrifuged at 21,000g for 10 min at 4°C to pellet the tissue. The supernatant containing metabolites was removed to a new Eppendorf tube and the metabolites were precipitated as described above. The tissue was then re-suspended in 1ml of 8M urea/50mM Tris-HCl, pH 8.0 and re-homogenized with the Polytron probe. The lysate was further homogenized by sonication as described above, cleared by centrifugation and processed for proteomic analysis as described above.

### DNA Extraction

50 ul of homogenized sample retained from the tissue or cell processing prior to tryptic digest was combined in an Eppendorf tube with 130 ul of nuclease-free water and 20 ul of 20 mg/ml Proteinase K, mixed briefly by vortexing (Thermo Fisher Scientific, Cat#AM2546) and incubated at 56°C for 1 hr. Following the digest, 200ul of phenol:chloroform:isoamyl alcohol (25:24:1) (Thermo Fisher Scientific, Cat#15593031) was added to the sample, the sample was vortexed thoroughly and centrifuged at 20,000g for 2 min at room temperature. 200 ul of the top (aqueous) phase was transferred to a new Eppendorf tube, combined with 2ul of 3M Sodium Acetate, pH 5.5 and 0.2 ul of 15mg/ml glycogen (Thermo Fisher Scientific, Cat#AM9516) and mixed by brief vortexing. 600 ul of 100% EtOH was added to the sample and the reaction was incubated at −20°C overnight. The sample was then centrifuged at 12,000g for 15 minutes at 4°C to pellet the DNA, the supernatant discarded, and the DNA pellet washed with 70% EtOH (centrifuged at 12,000g for 15 minutes at 4°C to pellet the DNA). The supernatant was discarded and the pellet air dried for 5 minutes and suspended in 20ul of 5 mM Tris-HCl pH 8.0, 1mM EDTA buffer.

### Quantitative Real-time PCR

Real-time PCR was performed using the Bio-Rad Laboratories Inc. CFX96 Thermal Cycler (Bio-Rad Laboratories Inc.) and iTaq SYBR Green Supermix (Bio-Rad, Cat#1725120). The following primers were used: ApoB forward (CACGTGGGCTCCAGCATT) and ApoB reverse (TCACCAGTCATTTCTGCCTTTG). The reaction was composed of ul iTAQ universal SYBR Green 2X master mix, 1ul (500nM) of forward primer, 1ul (500nM) of reverse primer, 1ul nuclease-free water and 2ul of the sample DNA (10ul total reaction), and the cycling conditions were 95°C 10 min, (95°C 15s, 60°C 15s, 72°C 15s) x 39 cycles. The calculations of average Cq values, SDs, and resulting expression ratios for each target gene was performed using the Bio-Rad Laboratories Inc. CFX Manager ver. 3.1.

### Microscopy

Phase-contrast images were captured with Keyence BZ-9000 fluorescence microscope (BIOREVO)

## QUANTIFICATION AND STATISTICAL ANAYSIS

### Statistical Analysis

One-way ANOVA with post-hoc Tukey HSD test was used to compare datasets with more than two groups. An unpaired t-test was used to analyze the rest of the datasets. Data are presented as mean ± SEM unless otherwise indicated. The number of biological and technical replicates is indicated in figure legends wherever appropriate. A p value of ≤0.05 was considered significant. GraphPad Prism 6, Excel 2016 and R Studio 3.5 software was used for analysis, Morpheus matrix visualization software used for metabolite heatmaps (https://software.broadinstitute.org/morpheus).

## Acknowledgements

The authors acknowledge the Cedars-Sinai Metabolism and Mitochondrial Research Core for respirometry studies.

## Author Contribution

A.S. and S.J.P. designed and directed the projects, A.O. and W.S. provided technical expertise and aided in sample preparation and instrument operation, A.A. assisted in data interpretation and guidance on simvastatin experiments, R.A.G., J.V.E. and S.J.P. provided funding acquisition for the project, S.J.P supervised the project. A.S. and S.J.P. wrote the manuscript.

