## Supplemental Figures for "MitoPlex: A Targeted Multiple Reaction Monitoring Assay for Quantification of a Curated Set of Mitochondrial Proteins"

**Supplemental Figure 1**

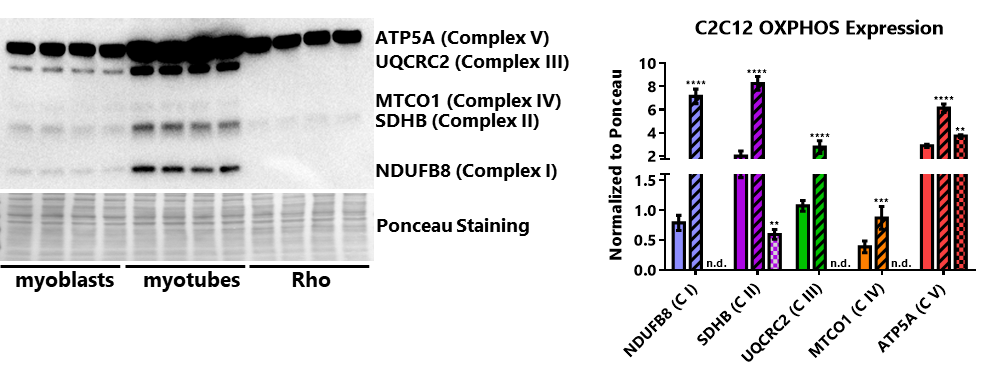

Western blot analysis of C2C12 myoblast, myotube and Rho lysates with the OXPHOS antibody cocktail and the quantification of the results, normalized to Ponceau staining (solid bars are myoblasts, stripes are myotubes and checkers are Rho). One-way analysis of variance (ANOVA) vs myoblasts, *p<0.05, **p<0.01, ***p<0.001; representative western blot is shown, values are means ±SD, n=4)

**Supplemental Figure 2**

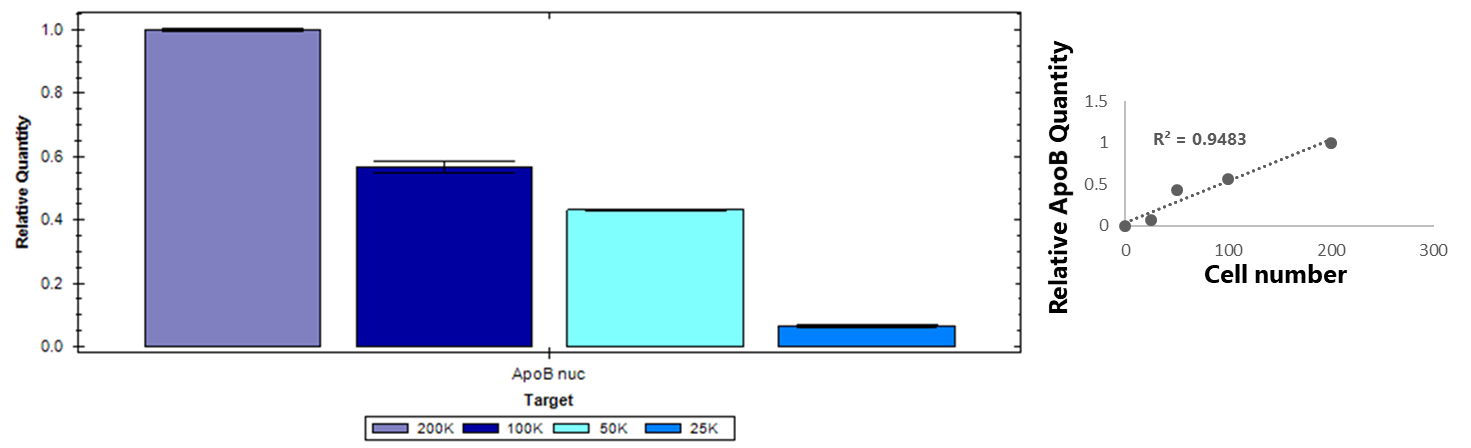

Relative ApoB signal as detected by quantitative PCR from DNA isolated from C2C12 myoblasts during the MitoPlex processing. Three technical replicates shown per sample.
